# Transcriptome sequencing of Hodgkin lymphoma Hodgkin and Reed-Sternberg cells reveals escape from NK cell recognition and an unfolded protein response

**DOI:** 10.1101/2025.08.05.667241

**Authors:** Mikhail Roshal, Isabella Y Kong, Wikum Dinalankara, Jonathan B. Reichel, Matthew Teater, Bhavneet Binder, Sakellarios Zairis, Joshua D. Brody, Sunita I. Park, Alexandra E. Kovach, Matthew J. Oberley, Megan S. Lim, Matthew J Barth, Olivier Elemento, Ari Melnick, Raul Rabadan, Amy Chadburn, Luigi Marchionni, Lisa Giulino-Roth, Ethel Cesarman

## Abstract

The mutational profile of classic Hodgkin lymphoma (cHL) overlaps with that of related B cell lymphomas, including primary mediastinal B cell lymphoma (PMBL), and yet these are different histologically and clinically. To discover the molecular features that distinguish cHL, we deployed flow cytometric cell sorting and low-input RNA sequencing to generate full transcriptome data from viable, isolated Hodgkin and Red-Sternberg (HRS) cells from eighteen primary tumors, alongside matched intra-tumoral non-neoplastic B cells and four cell lines. Comparison of HRS cells to normal cellular subsets revealed evidence of abortive plasma cell differentiation, with an unfolded protein response signature, shared with plasma cell neoplasms, but not other B-cell lymphoma types. Comparison of cHL to PMBL revealed similarities but also key differences in B cell differentiation programs accompanied by upregulation of genes involved in microtubule cytoskeleton organization in cHL, which may be related to the unique multinucleated nature of HRS cells. In HRS cells, we also observed a downregulation of SLAM family receptors, which are crucial for NK cell activation, providing a potential mechanism for immune evasion from NK-mediated killing.

**STATEMENT OF SIGNIFICANCE:** This study defines a unique transcriptional program in classic Hodgkin Lymphoma (cHL) marked by oncogenic signaling, chromatin integrity, DNA repair, and immune escape including loss of NK-activating receptors. HRS cells resemble plasma cells and have an unfolded protein response signature, which distinguishes them from diffuse large and primary mediastinal B cell lymphomas.

## INTRODUCTION

The malignant cells of classic Hodgkin Lymphoma (cHL), known as Hodgkin and Reed-Sternberg (HRS) cells, are embedded within a dense infiltrate of benign immune and stromal cells. Consequently, >99% of the bulk genomic material of a cHL lymph node is typically derived from non-neoplastic cells (1). As a result, obtaining high-quality tumor-derived nucleic acids, free from significant background contamination, necessitates the physical separation and concentration of HRS cells. Efforts to fully characterize the molecular and genomic landscape of primary HRS cells have been hindered by the challenges associated with isolating sufficient quantitates of pure HRS cell material to generate robust data.

Several studies have used cultured cHL cell lines as a surrogate for primary HRS cells. For example, microarray studies have shown that cHL cell lines clustered distinctly from normal B cells and B cell non-Hodgkin lymphomas (NHLs) (2). RNA sequencing of two cell lines uncovered a highly expressed gene fusion involving CIITA (Class II major histocompatibility complex transactivator, or MHC2TA), which was found to be recurrent in approximately one-third of primary mediastinal B-cell lymphoma (PMBL) and 15% of cHL (3). Other studies have used microdissected HRS cells for comparative genomic hybridization to identify chromosomal imbalances (4). In 2012, two independent studies utilized microarray technology to analyze microdissected HRS cells from primary tumors, revealing that primary HRS cells differ from cHL cell lines, other B-cell lymphomas, and normal B cells(5,6). Recently, Weniger et al. demonstrated that the transcriptome of microdissected HRS cells closely resembles that of CD30+ B cells, particularly extrafollicular CD30+ B cells, possibly due to shared CD30-mediated signaling pathways (7). Notably, a key difference between CD30+ B cells and HRS cells was the downregulation of genes involved in cytokinesis and genomic instability, which may underlie the frequent multinuclearity and genomic instability observed in cHL (7).

We previously reported the first whole exome sequencing of primary HRS cells to identify somatic mutations, small indels, and copy number alterations, utilizing a combined approach of cell sorting and optimized low-input sequencing across ten primary cases and two cell lines (8). Subsequent exome sequencing by Wienand *et al*. confirmed many of the genetic alterations identified in our study, offering a more accurate frequency assessment due to a larger cohort (9). Other techniques, such as FACS sorting of nuclei (10) and analysis of circulating tumor DNA (11–13), have also been employed to investigate the cHL genetic landscape. Our group further extended this work by reporting the first full genome sequences of purified HRS cells, as well as exome sequencing in a larger cohort, providing a map of genomic alterations and insights into evolutionary timing in both pediatric and adult cHL (14). Our findings suggested that while mutations and genomic alterations in cHL occur both pre- and during germinal center entry, the terminal HRS cells likely do not complete the germinal center reaction with transition from dark to light zone and back, given the absence of an SBS9 mutational signature (potentially mediated by DNA polymerase eta) that likely occurs after AID-induced somatic hypermutation, but is present in other lymphomas of GC origin (14,15).

Characterization of HRS DNA, including from cell lines, primary tissue samples and circulating tumor DNA, has revealed a spectrum of mutations that overlap with those of diffuse large B cell lymphomas (DLBCL) and primary mediastinal B cell lymphomas (PMBL) (16–22). We hypothesized that the HRS transcriptome would explain at least some of the major differences between these clearly distinct malignancies. Here, we present the first whole transcriptome sequencing of primary HRS cells from eighteen primary cases (seventeen with matched intra-tumoral B cells) and four cell lines. Our analysis reveals that HRS cells exhibit abortive plasma cell differentiation, as evidenced by an increased unfolded protein response signature, alongside novel alterations in pathways involved in oncogenesis, cell division, and immune evasion.

## RESULTS

### Distinct gene expression profiles of HRS cells compared to intra-tumoral B cells

To examine the transcriptome of HRS cells, we conducted bulk-RNA sequencing on primary HRS cells isolated from 18 primary cases of cHL, comparing them to intra-tumoral non-neoplastic B cells (available in 17 cases) and four HL cell lines (KMH2, HDLM2, L1236, and L428) (**Supplementary Table S1**). Cases were numbered 2-19, where 2 through 10 match those reported for exome sequencing (8). Principal component analysis (PCA) and unsupervised clustering analysis revealed two distinct clusters that segregated intra-tumoral B cells from both HRS cells and HL cell lines (**Figure 1A-B**). Volcano plots are shown for differentially expressed genes between HRS cells and intratumoral B cell, cHL cell lines and intratumoral B cells and HRS cells and cHL cell lines (**Figure 1C-E**). Comparative analysis between HRS cells and intra-tumoral B cells revealed 2144 upregulated genes (logFC≥2; adjp≤0.01) and 2451 downregulated genes (logFC≥-2; adjp≤0.01) (**Supplementary Datatable S1 and S2**). Based on these results, we defined the top 200 upregulated and top 200 downregulated genes (**Supplementary Datatable S3)**, which are referred to herein as “the cHL signature”. To determine if our RNAseq data were aligned with prior reports, we compared our samples with previously published gene expression profiles of microdissected HRS cells from 29 cases and 5 cell lines, generated using Affymetrix arrays (23). After filtering for significance, both technologies and analyses yielded correlated gene-level fold-change values across datasets, with 20 genes being differentially expressed in opposite directions (Pearsons’s coefficient 0.86; Spearman’s correlation 0.73; **Supplementary Figure S1**, **Supplementary Table S2**). While trends for most genes confirm this prior study, the current RNAseq data provides more sensitivity and specificity and reveals expression of genes not present in the microarray, allowing for precise pathway analysis and virus identification.

**Figure 1:**
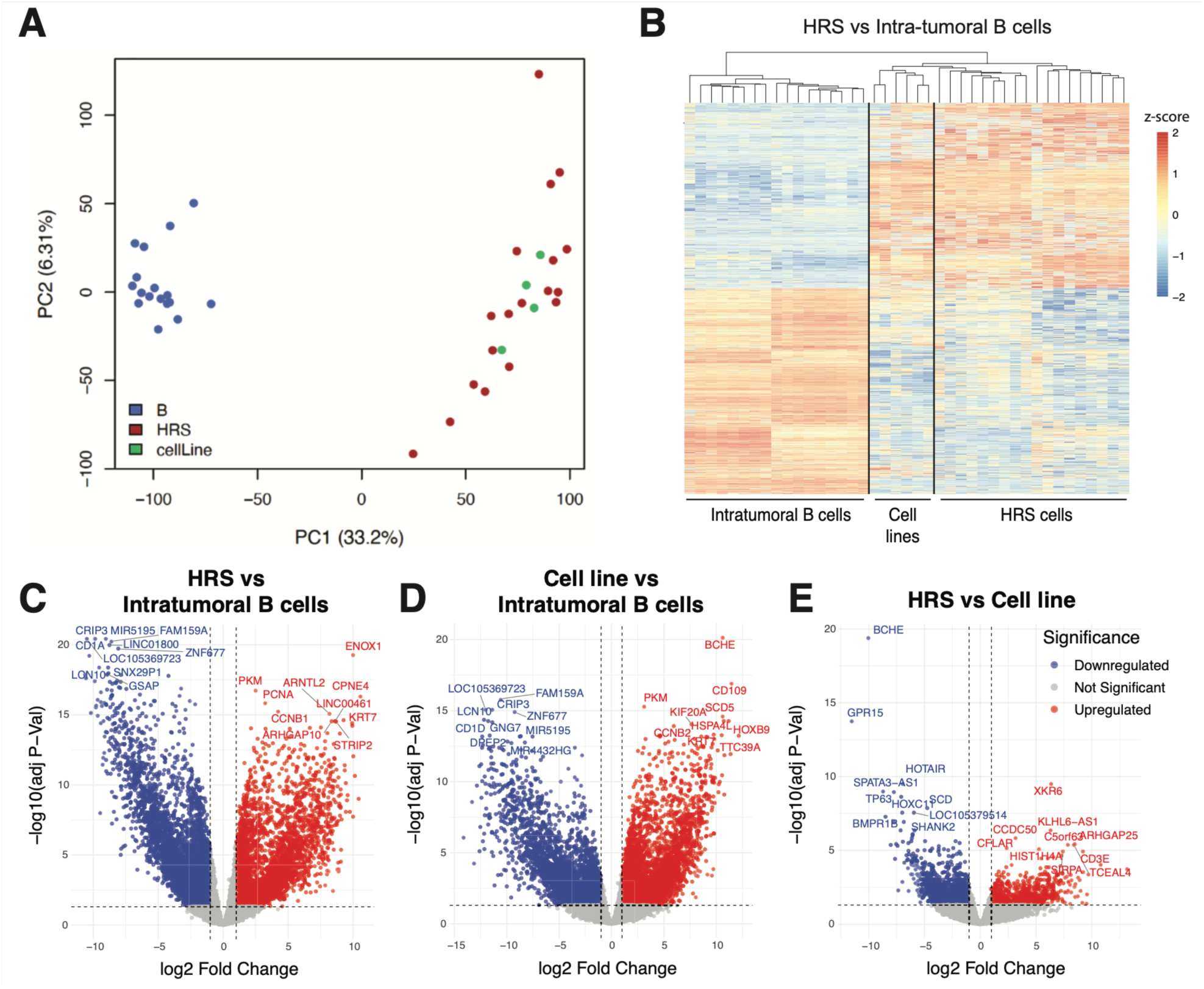
Distinct gene expression profiles of HRS cells compared to intra-tumoral B cells. (**A**) Principal component analysis (PCA) of gene expression profiles from HRS cell populations from 18 primary cHL cases (red) and intratumor B cells from 17 primary cHL cases (blue) and four HL cell lines (KMH2, HDLM2, L1236, and L428) (green). (**B**) Unsupervised hierarchical clustering of the differentially expressed genes between HRS and intra-tumoral B cells. Colors in **B** represents relative expression: red indicates higher and blue indicates lower expression relative to the gene’s mean. Volcano plot for genes that are differentially regulated between (**C**) HRS and intratumoral B cells, (**D**) HL cell lines with intratumoral B cells and (**E**) HRS cells and HL cell lines.

### Increase in NFKB and E2F family signaling in HRS cells

We first examined the genes that are significantly upregulated in HRS cells compared to intra-tumoral B cells. Gene ontology analysis of upregulated genes in HRS cells (logFC≥2; adj-pvalue≤0.01) revealed genes involved in processes such as the mitotic cell cycle, tube morphogenesis, and cell development (**Figure 2A**). This validates previous reports suggesting that the multinucleated HRS cells exhibit an abortive mitotic cycle, accompanied by persistent microtubule bonding due to incomplete cytokinesis, which may explain the multinucleated nature of these cells (24,25). To further explore the pathways upregulated in HRS cells relative to intra-tumoral B cells, we performed Gene Set Enrichment Analysis (GSEA). The GSEA results indicated an enrichment of genes associated processes that are known to be dysregulated in HL including: G2M checkpoint, IL2 STAT5 signaling, MYC targets, TNFα signaling via NFKB, and “inflammatory response” (**Figure 2B**). HRS cells are characterized by aberrant activation in several signaling pathways including those resulting from recurrent mutations in components of the nuclear factor-kB (NF-KB) pathway and the JAK/STAT pathways. Here, we confirmed that HRS cells have high expression of *NFKB* transcript levels and enrichment in the NF-κB signaling pathway (**Figure 2C**). Transcription factor target over-representation analysis for genes that are upregulated in HRS in comparison to intra-tumoral B cells (adj-p≤0.001, logFC≥2.32) also identified NFKB2 as one of the transcription factor activities to be upregulated in HRS cells (**Figure 2D**).

**Figure 2.**
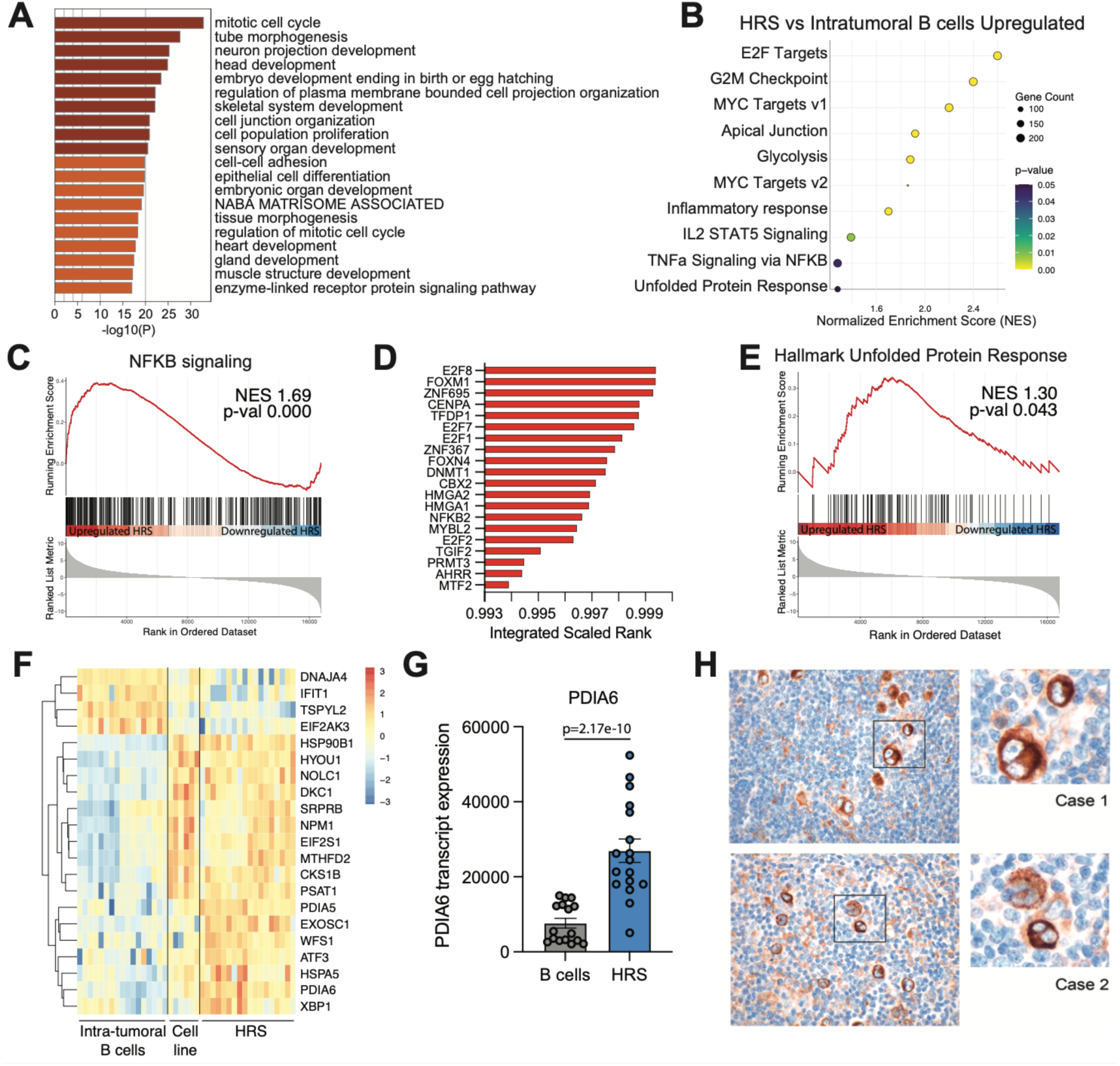
Upregulation of the unfolded protein response (UPR) in HRS cells. **(A)** Gene ontology analysis for genes that are significantly upregulated in HRS in comparison to intra-tumoral B cells (logFC≥2; adj-pval≤0.05). (**B**) Ten Hallmark pathways identified by GSEA for genes that are differentially regulated between HRS and intra-tumoral B cells. (**C**) GSEA of NFKB signaling pathway. (**D**) Transcription factor target over-representation analysis was performed using ChIP-X Enrichment Analysis version 3 (ChEA3) for genes that are upregulated in HRS in comparison to intra-tumoral B cells (adj-p≤0.001, logFC≥2.32). (**E**) GSEA for the unfolded protein response (Broad Institute MSigDB: HALLMARK_UNFOLDED_PROTEIN_RESPONSE) comparing HRS cells to intra-tumoral B cells. (**F**) Heatmap of differentially regulated UPR genes between HRS and intra-tumoral B cells (adjp≤0.05; logFC ≥ 1 or logFC ≤ −1) based on UPR gene signature in **B**. (**G**) Transcript expression of PDIA6 in intra-tumoral B cells (gray) and HRS cells (blue), with adjusted-p value determined using limma-voom. (**H**) Immunohistochemistry for PDIA6 in two representative cHL cases. Original magnification: 60X. Colors in **F** represents relative expression: red indicates higher and blue indicates lower expression relative to the gene’s mean.

In addition to NFKB2, we also identified several E2F family proteins (E2F8, E2F7, E2F1 and E2F2) as transcription factors predicted to be upregulated in HRS cells (**Figure 2D**). E2F family members are involved in many cellular processes and have been shown to play a role in cancer development and progression. Members of the E2F family, such as E2F1 and E2F6 have been shown to be dysregulated in cHL and are correlated with proliferation and apoptosis (26,27). Additionally, our analysis also identified several transcription factors known to be associated with lymphomas and other malignancies, including FOXM1 and CENPA. We also identified several zinc finger proteins, including ZNF695 and ZNF367, that have been associated with B cell acute lymphoblastic leukemia (B-ALL) and DLBCL respectively (28,29).

### HRS cells exhibit an unfolded protein response

GSEA also revealed a significant enrichment of pathways involved in biological processes not previously described in cHL, including the unfolded protein response (UPR) (**Figure 2B, 2E-F**). To assess whether this UPR signature is specific to HRS cells, we evaluated its expression in DLBCL and primary mediastinal B-cell lymphoma (PMBL), a subtype of NHL with molecular features similar to cHL (19,21,30), using previously reported gene signatures for these diseases (18–21). Multiple myeloma (MM), a plasma cell malignancy with well-characterized UPR activation, was included as a positive control. As expected, MM samples exhibited elevated UPR signature and increased expression of UPR-related genes (**Supplementary Figure S2A-B**). In contrast, we did not observe a consistent increase in UPR signature expression in PMBL or in either the activated B-cell (ABC) or germinal center B-cell (GCB) subtypes of DLBCL (**Supplementary Figure S2B-F**). While a subset of ABC-DLBCLs do have partial plasmacytic differentiation (31) and a degree of UPR (**Supplementary Figure 2E**), the consistent UPR activation is a distinct feature of HRS cells in cHL.

Among the top upregulated UPR genes, we identified PDIA6, a disulfide isomerase localized in the endoplasmic reticulum (ER) that plays a crucial role in protein folding and preventing aggregation by catalyzing the formation and breakage of disulfide bonds during protein folding (**Figure 2F-G**). Immunohistochemistry (IHC) using PDIA6 antibodies in cHL cases 2-10, as well as in a tissue microarray (TMA) containing an additional 16 cases, demonstrated strong immunoreactivity in HRS cells compared to background lymphoid cells in 100% of the cases (**Figure 2H**). The intensity of PDIA6 staining was similar to or greater than that observed in residual plasma cells, which are characterized by an UPR. Apart from plasma cells, which are easily recognized morphologically, PDIA6 staining was highly specific for HRS cells, distinguishing them from other cell types, and therefore may serve as a valuable diagnostic marker.

### Downregulation of B-cell program and NK cell recognition pathways in HRS cells

We next examined the genes that are downregulated in HRS in comparison to intra-tumoral B cells. As expected, GSEA revealed a downregulation of immune regulatory processes such as B cell receptor signaling, antigen processing and presentation, and B cell activation, all of which are known to be suppressed in HRS cells (**Figure 3A**). This observation aligns with the gene ontology (GO) analysis of the downregulated genes in HRS (logFC≤-2; adj-pval≤0.01), which also indicated downregulation of these pathways (**Supplementary Figure S3A-C**).

**Figure 3.**
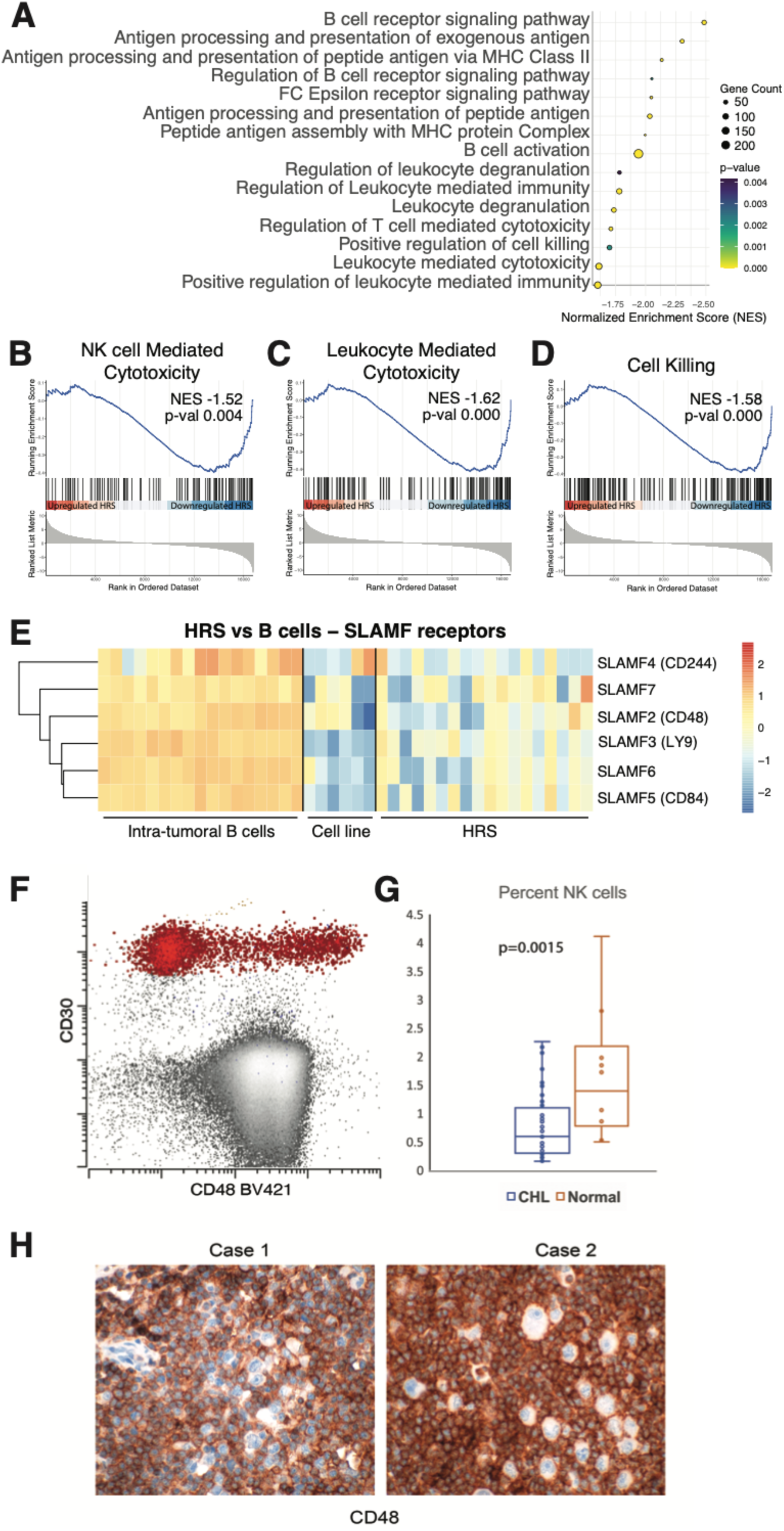
Downregulation of NK cell recognition pathways in HRS cells. (**A**) 15 downregulated biological processes identified between HRS and intra-tumoral B cells. GSEA for (**B**) KEGG Natural killer cell mediated cytotoxicity, (**C)** GOBP Leukocyte mediated cytotoxicity and (**D**) GOBP Cell killing pathways comparing HRS to intra-tumoral B cells. NES represents normalized enrichment score. (**E**) Heatmap displaying differential expression of SLAMF receptors in HRS vs intra-tumoral B cells (adjp≤0.001, logFC≤-1). **(F)** Flow cytometry analysis of CD48 expression on CD30+ HRS cells (red) compared to CD30-cells in a primary cHL specimen. **(G)** Flow cytometry quantification of NK cell proportions in cHL (N=48) and normal lymph nodes (N=10) samples. **(H)** Immunohistochemistry for CD48 across two representative cHL cases. Original magnification: 60X. Colors in **E** represents relative expression: red indicates higher and blue indicates lower expression relative to the gene’s mean.

The downregulation of B cell activation and B cell receptor signaling is further supported by the transcription factor target over-representation analysis, which indicated a reduction in the activity of key B cell transcription factors such as IRF8, Pax5 and Pou2f2 (**Supplementary Figure S3D**). Notably, we also observed a downregulation of SP140, an “epigenetic reader” that primarily represses immune-related genes through NKFB inhibition (32). Inactivating SP140 mutations have been found in chronic lymphocytic leukemia (33) and multiple myeloma (34), but overexpression has been reported in the ABC subtype of DLBCL (35). In Burkitt lymphoma, it was shown to bind PRC2 and NuRD complexes directing H3K27me3 on target genes (34), but its downregulation in cHL had not been previously established.

GSEA also identified several pathways related to leukocyte activation and degranulation, which have not been previously associated with cHL (**Figure 3A**). To further investigate the specific leukocyte subsets contributing to these signals, we examined cell type-specific pathways and found that natural killer (NK) cell-mediated cytotoxicity emerged as significantly enriched in the analysis. Further comparisons revealed that pathways related to NK cell mediated cytotoxicity, leukocyte-mediated toxicity and cell killing were downregulated in HRS cells in comparison to intra-tumoral B cells (**Figure 3B-D**).

One of the key regulators of NK cell cytotoxicity is the signaling lymphocytic activation molecule family (SLAMF) receptors. Thus, we examined the expression of SLAMF receptors on HRS cells. Our results showed that six out of nine SLAM family activating receptors - *SLAMF2* (*CD48*), *SLAMF3* (*LY9*), *SLAMF4* (*CD244*), *SLAMF5* (*CD84*), *SLAMF6* and *SLAMF7* – were downregulated on HRS cells (**Figure 3E**), suggesting that the loss of NK cell-activating receptors may contribute to immune evasion by HRS cells. We further validated this finding using flow cytometry, confirming the downregulation of CD48 (SLAMF2) in HRS cells from both cell lines and five primary cases (**Figure 3F)**. Consistent with the reduced role of NK cells in tumor immune control, we found a significantly reduced proportion of NK cells in cHL tumors (median 0.6% vs. 1.4%, p<0.01) compared to reactive lymph nodes, as assessed by flow cytometry (**Figure 3G**). Immunohistochemistry confirmed the loss of CD48 in every CHL case examined, including the nine sequenced cases (cases 2-10) and 15 additional specimens (**Figure 3H**).

### Comparative transcriptomic analysis of cHL reveals key differences between cHL and PMBL and similarities to CD30+ B cells, and both normal and malignant plasma cells

Exome-wide genome studies have demonstrated substantial mutational overlap between PMBL and cHL. To investigate the transcriptional differences that distinguish these entities, we compared the PMBL gene signature defined by Savage *et al.* (20) to gene expression profiles from intratumoral B cells, cHL cell lines and purified HRS cells, identifying a subset of differentially regulated genes (**Figure 4A)**. Notably, *TNFRFS17* (encoding BCMA), typically expressed in plasma cells, and *LMO2,* associated with germinal center B cells, were upregulated in PMBL but uniformly absent in cHL. Immunohistochemistry on an extended cohort confirmed BCMA and LMO2 expression in PMBL cases (BCMA 45/49, LMO2 50/50), but not in HRS cells of cHL biopsies, (BCMA 0/17; LMO2 0/13) (**Figure 4B).** To further delineate shared and distinct transcriptional features between cHL and PMBL, we assessed the expression of differentially expressed genes (DEGs) between HRS cells and intratumoral B cells from PMBL samples, using data from Mottok et al. (19) (**Figure 4C and Supplementary Data**t**able S6**) (19). We identified 1072 genes upregulated in HRS cells that are also highly expressed in PMBL. GO analysis of these genes indicated enrichment in pathways related to cell cycle regulation and oncogenic signaling (**Figure 4D**). Conversely, 1412 genes downregulated in HRS cells and lowly expressed in PMBL were associated with immune and inflammatory response pathways (**Figure 4E**). To investigate transcriptional differences between cHL and PMBL, we performed GO analysis on genes whose expression was discordant between cHL and PMBL. The 1434 genes upregulated in HRS cells but downregulated in PMBL were enriched for neuronal processes and microtubule cytoskeleton organization. In contrast, 1412 genes downregulated in HRS cells and highly expressed in PMBL were involved in B cell receptor (BCR) signaling and immune activation. These findings are consistent with disordered cytokinesis and the loss of B cell surface markers in HRS cells, both of which are features of cHL that are not observed in PMBL.

**Figure 4.**
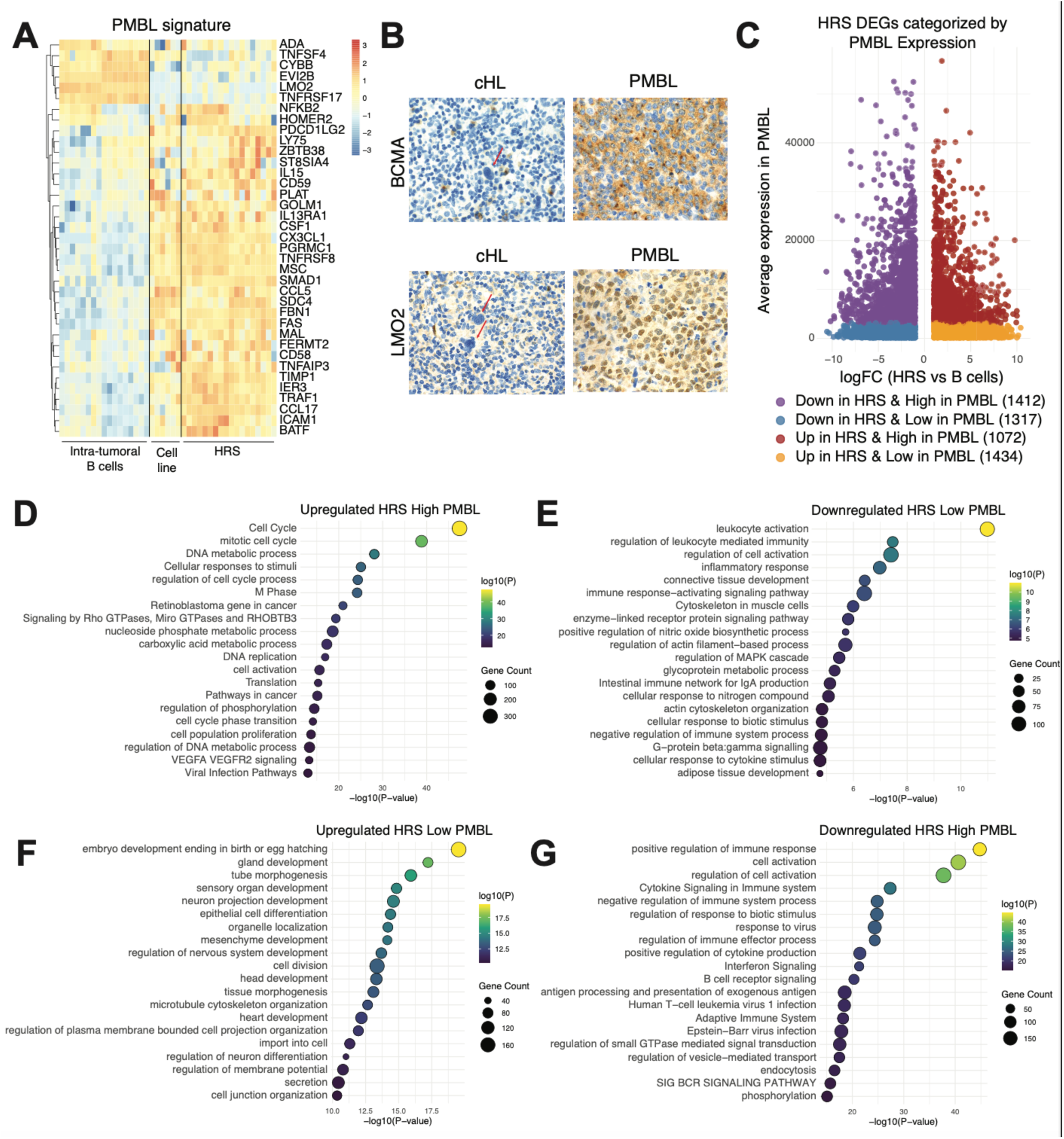
Comparative transcriptomic analysis of HRS cells and primary mediastinal B-cell lymphoma (PMBL). (**A**) Heatmap showing the expression of PMBL gene signature defined by Savage *et al.* (20) in intratumoral B cells, HL cell line and HRS cells. (**B**) IHC for BCMA (cytoplasmic, top panels) and LMO2 (nuclear, bottom panels) for cHL and PMBL tumors. HRS cells are depicted by red arrows. Cases were those included in a tissue microarray and only considered evaluable among the cHL cases if internal non-tumoral positive control cells were present. Original magnification was 60x. (**C**) Scatter plot of differentially expressed genes between HRS cells and intratumoral B cells (adjp-val≤0.05; logFC≥1 or logFC≤-1) compared to gene expression in PMBL (19). Gene ontology analysis was performed on genes that are (**D**) upregulated in HRS cells vs intratumoral B cells and highly expressed in PMBL, (**E**) downregulated in HRS cells vs intratumoral B cells and lowly expressed in PMBL, (**F**) upregulated in HRS cells vs intratumoral B cells and lowly expressed in PBCL, and (**G**) downregulated in HRS cells vs intratumoral B cells and highly expressed in PMBL. Gene expression data for PMCL (panels **C-G**) were obtained from Mottok et al (19). Colors in **A** represents relative expression to the gene’s mean.

Previous studies have suggested that HRS cells may originate from CD30^+^ cells within the germinal center. To investigate this, we first examined whether the CD30^+^ signature (7) is enriched in our cHL signature. Indeed, we found that genes upregulated in CD30^+^ cells were significantly enriched (NES:2.21; p-val:0.000) (**Figure S4A**). To explore the possibility that HRS cells might derive from other non-malignant B cell subsets, we examined the cHL signature genes across various B cell subsets. The HRS cell transcriptional profile was closer to that of centroblasts than centrocytes, consistent with a closer relationship to dark zone B cells. Notably, the expression patterns of these genes in HRS cells most closely resembled those found in plasma cell lineages, including bone marrow plasma cells and, but to a lesser extent tonsillar plasma cells (**Figure 5A-B**). Further analysis to assess signatures assigned by Cibersortx revealed that, compared to intra-tumoral B cells, HRS cells exhibited a transcriptomic signature more closely resembling bone marrow plasma cells and GCBs (**Figure 5C-D, S4B**).

**Figure 5.**
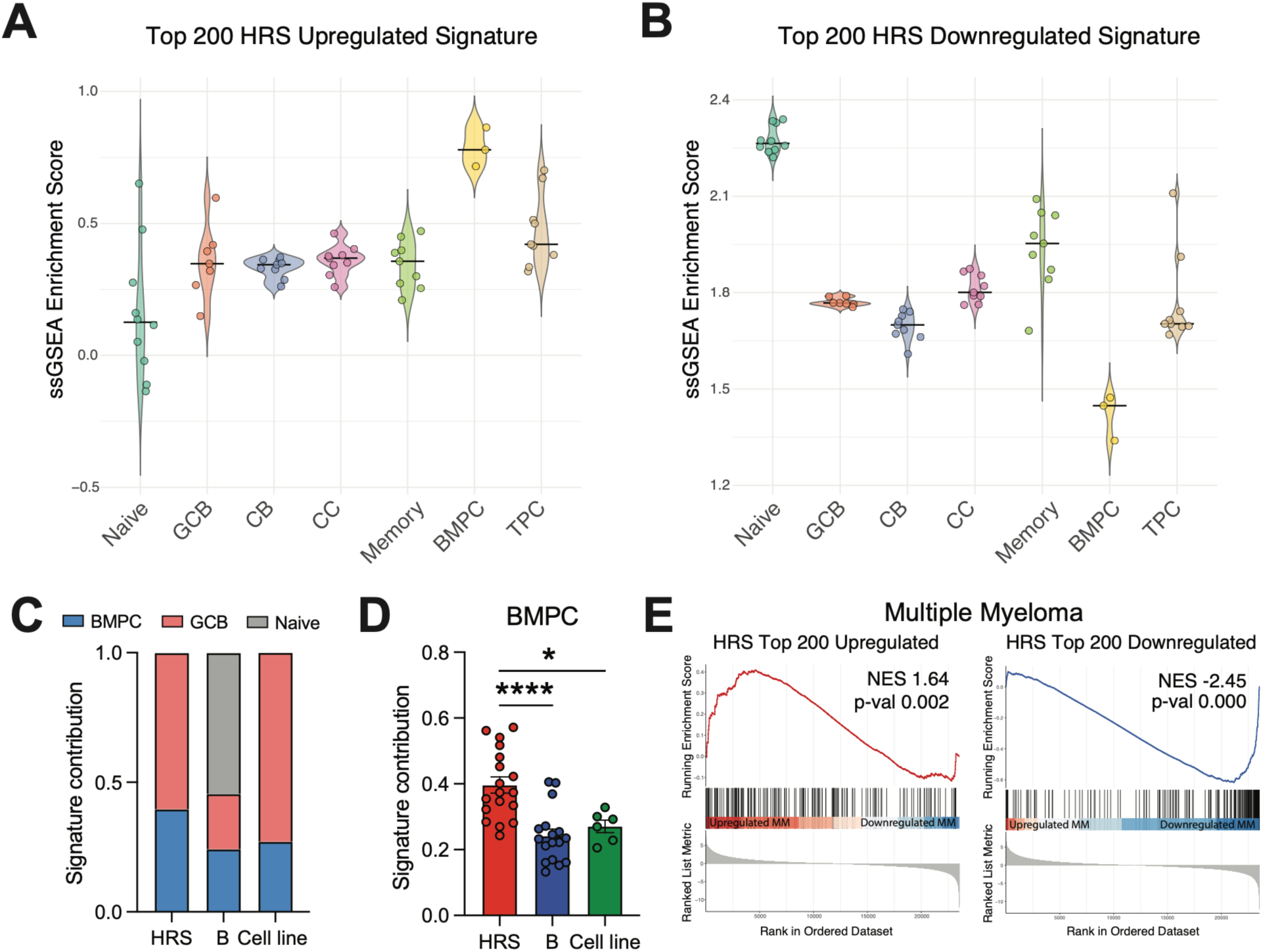
Comparative transcriptomic analysis of HRS cells suggests a germinal center-derived origin and similarities to plasma cell lineages. Enrichment scores of the (**A**) top 200 upregulated and (**B**) top 200 downregulated genes in HRS vs intra-tumoral B cells across different B cell subset (BMPC: bone marrow plasma cells; CB: centroblasts; CC: centrocytes; GCB: germinal center B cells; Memory: memory B cells; Naïve: naïve B cells; TPC: tonsil plasma cells). (**C**) Cibersortx analysis estimating the abundance of bone marrow plasma cells (BMPC), germinal center B cells (GCB) and naïve B cells signature in HRS cells, intra-tumoral B cells and HL cell lines. (**D**) Estimated abundance of BMPC signature in HRS cells, intra-tumoral B cells and HL cell lines. (**E**) GSEA for top 200 upregulated and downregulated genes in HRS vs intra-tumoral B cells in multiple myeloma (MM) samples. NES in **E** represents normalized enrichment score. Colors in **A-B** represents relative expression to the gene’s mean.

Given the overlap between HRS cells and plasma cells, we next examined the expression of the cHL signature in multiple myeloma (MM). We found that the genes upregulated in the cHL signature were positively enriched MM vs. GCBs (NES: 1.64; p=0.002), and conversely those negatively regulated were reduced in MM (NES: −2.45, p=0.000) (**Figure 5E**).

### Virus discovery

We performed a search for known and possible novel infectious agents using the Pandora pipeline in cases 2-9 and the four cell lines (36), and we followed up and tested all the cases using the Virdetect pipeline (37). We detected Epstein-Barr virus (EBV) in a single sample (case 8), recovering 19 contigs above our cutoff of 500bp, the longest of which was 1829bp. Case 8 was the only case among those evaluated known to be positive for EBV, and EBV expression profiling showed most abundant expression of genes *LMP1* and *LMP2*, consistent with its known latency II pattern (Supplementary Figure S5). The only other virus that was found was bovine viral diarrheal virus (BVDV), which was present in the HRS cells of cases 2, 3, and 4, and B cells from cases 3 and 7. BVDV is a common contaminant of fetal calf serum (**Supplementary Datatable S5**). To determine whether previously undiscovered viruses were present, we performed alignments of all contigs assembled from non-human reads that did not match known sequences using low-stringency alignments. No credible matches were identified, suggesting that if new viruses exist in cHL, they are very divergent to existing sequences in databases or they occur only in a small proportion of cHL.

## DISCUSSION

We present the first RNA sequencing dataset of HRS cells isolated from primary cases of cHL. Our transcriptomic analysis reveals that cHL shares greater similarity with plasma cells than with mature B cells and shares features with plasma cell malignancies, such as multiple myeloma (MM). While prior studies have established a germinal center B cell origin for cHL, supported by the presence of ongoing somatic hypermutation (38–41), our findings provide molecular evidence for partial plasmacytic differentiation, in line with prior immunohistochemical observations of abortive plasma cell features in HRS cells (42,43).

Despite genomic and transcriptomic overlaps with DLBCL and PMBL, HRS cells exhibit a distinct gene expression profile enriched for a plasma cell-related signature, which is driven at least in part by UPR. This profile is absent in most DLBCL and PMBL cases, but present in MM, further supporting a unique and aberrant differentiation state. Notably, HRS cells show greater transcriptional similarity to bone marrow plasma cells than to tonsil plasma cells; this is consistent with bone marrow plasma cells arising from cells that have transited through a germinal center reaction (44). We previously reported an unusual dissociation between the canonical AID signature and SBS9 signature in cHL, where there is AID but not SBS9. Although SBS9 was initially attributed to non-canonical AID activity, recent studies suggest it is more closely linked to replicative stress in GC B cells (15). While such dissociation has been observed in normal B cells (15), it has not been reported in B cell lymphomas outside of cHL. Interestingly, a similar pattern occurs in a subset of MM cases with MAF rearrangement, suggesting that cHL and specific MM subsets may share mechanisms governing GC exit. Downregulation of FOXO1, as previously shown by our group and others, and confirmed in this study, likely contributes to defective dark zone programs in HRS cells (45). Together, these findings support a model in which HRS cells aberrantly exit the germinal center through an early abortive plasma cell differentiation program further compromised by failure to express immunoglobulin.

Our comparative transcriptomic analysis highlights the similarity between cHL and PMBL. The overlap in upregulated genes related to cell cycle and oncogenic pathways suggests common mechanisms driving proliferation in both malignancies. Conversely, the downregulation of immune-related genes, particularly those involved in BCR signaling, reinforces the idea that HRS cells undergo a profound loss of B cell identity, unlike PMBL. While PMBL tumors co-express LMO2, a GCB cell marker, and BCMA, a plasma cell marker, HRS cells lack both, indicating that each malignancy follows an aberrant but distinct path of B cell differentiation. Interestingly, the unique enrichment of neuronal and cytoskeletal gene programs in HRS cells may reflect lineage infidelity or aberrant differentiation processes not observed in other B cell malignancies.

A hallmark of terminal plasma cell differentiation is activation of the unfolded protein response (UPR), which we observed in both HRS cells and MM but not to a comparable extent in most DLBCL or PMBL cases. However, plasma cell differentiation in cHL appears incomplete. In contrast with plasmablasts, HRS cells typically do not express CD138 (46), although exceptions have been reported (47). While IRF4 (MUM1) is uniformly overexpressed, *PRDM1* (BLIMP1), a key regulator of plasma cell maturation, shows relatively low expression levels, consistent with previous studies (5,42,48). This imbalance likely contributes to the absence of immunoglobulin production in HRS cells, despite robust UPR activation, as evidenced by consistently high expression of PDIA6 and other UPR components. While prior studies have reported overexpression of UPR components such as *XBP-1* and *ATF6* in cHL (49), our data extend these findings by demonstrating a robust activation of the full UPR transcriptional program. In MM, UPR activation is partially driven by PRDM1 (50). The relatively low PRDM1 levels in cHL, potentially due to *FOXO1* downregulation (48), suggests an alternative mechanism of UPR induction.

Our data also provide an integrated view of immune evasion mechanisms employed by HRS cells that may underlie the absence of effective anti-tumor T cell or innate immune responses in cHL. HRS cells exhibit impaired antigen presentation due to *B2M* mutations, overexpression of immune checkpoint ligands (*PDL1*, *PDL2*, *TIM3)*, and secretion of immunosuppressive cytokine. Notably, we report for the first time the absence of SLAM family NK cell activating receptors on HRS cells (e.g. CD48), which may contribute to their evasion of NK cell-mediated cytotoxicity, especially given concurrent MHC class I downregulation. Furthermore, we observed a marked reduction in NK cell infiltration in cHL tumors compared to reactive lymph nodes, suggesting spatial exclusion of NK cells as an additional immune evasion strategy.

While cHL has long been hypothesized to harbor a yet undiscovered viral pathogen in addition to EBV (51), our transcriptomic analysis did not reveal any novel viral sequences. EBV transcripts were detected only in EBV-positive cases, consistent with prior reports (52–54). Our previous whole-exome sequencing (8), along with other studies (55), showed that *TNFAIP3 (A20*), a negative regulator of NFkB, is frequently mutated in EBV-negative cHL but typically intact in EBV-positive cases. However, this correlation is not absolute, as demonstrated by a larger study (14). EBV-positive tumors exhibit lower overall mutational burden, supporting a model in which EBV-driven activation of NFkB via latent membrane proteins LMP1 and LMP2 substitutes for somatic mutations that drive similar pathways in EBV-negative cHL.

## METHODS

### Tissue specimens

Eighteen cases underwent RNA sequencing, which were among those previously reported for exome sequencing (8) and numbered accordingly 2-10 to correspond with previously published cases; cases 11-16 were published by Maura *et al*. (14) (Supplementary Table S1). Cases were from the Department of Pathology and Laboratory Medicine at Weill Cornell Medical College (WCMC, case 2-9), Mount Sinai Medical Center (case 10), Children’s Hospital of Los Angeles, Children’s Healthcare of Atlanta, Children’s Hospital of Philadelphia, Roswell Park Cancer Center, and Memorial Sloan Kettering Cancer Center. All cases were deidentified and obtained with institutional IRB approval. Case 1 from the prior study(8) had no residual material, so it was excluded from RNA analysis, but the nomenclature has been maintained to allow for comparative mining of DNA and RNA sequencing data. Seventeen of these cases (2–18) had paired intra-tumoral B cell sequenced, and one (case 19) did not, and data from these specimens was designated as HRS cells or to B cells respectively for each case. Group size was limited by availability of fresh frozen viable cell suspensions and recovery of HRS cells, which determined the final cohort of 19. Specimens were mechanically dissociated and cryopreserved as viable cell suspensions following excisional or needle core lymph node biopsy. Formalin-fixed paraffin-embedded (FFPE) cHL biopsies were used for immunohistochemical validation.

### Immunohistochemistry

IHC was performed on tissue sections (cases 2-10), a cHL microarray containing 16 additional cases and a PMBL TMA containing 50 cases. Immunohistochemical staining of CD48 (1:100, rabbit monoclonal, clone EPR4108, Abcam, RRID:AB_10703015), PDIA6 (1:1000, Rabbit polyclonal, Proteintech, RRID:AB_10805765), BCMA (1:200, mouse monoclonal, clone D-6, Santa Cruz Biotechnology) and LMO2 (1:2000, mouse monoclonal, clone 1A9-1, Santa Cruz Biotechnology, RRID:AB_2136572) was accomplished using the Bond III auto stainer (Leica Microsystems, Illinois, USA). FFPE tissue sections were first baked and deparaffinized. Antigens were then retrieved by heating the slides at 99-100°C in Bond Epitope Retrieval Solution 2 for 20 minutes for CD48 and 30 minutes for BCMA and LMO2, and in Bond Epitope Retrieval Solution 1 for 30 minutes for PDIA6. Sections were then incubated sequentially with the endogenous peroxidase block for 5 minutes, primary antibody for 15 minutes (PDIA6 and CD48) or 30 minutes (BCMA and LMO2), polymer (equivalent to tertiary antibody) for 8 minutes, diaminobenzidine (DAB) for 10 minutes and hematoxylin for 5 minutes (Bond Polymer Refine Detection, Leica Microsystems). The stained slides were then dehydrated in 100% ethanol and mounted by using LeicaCV5030 Glass Coverslipper (Leica Microsystems).

### Cell Sorting

HRS cells and non-neoplastic intra-tumoral B cells were separated from inflammatory background cells in lymphoid tissue using FACS, as previously reported (8,56). To separate HRS cells and non-neoplastic intratumor B cells from inflammatory background cells in lymphoid tissue, we used the method previously reported (48,56,57). Briefly, cell suspensions from cHL primary tumor cases were rapidly defrosted in a 37C water bath and washed with RPMI 1640/20% fetal bovine serum solution containing DNase A. Cells were then stained for 15 minutes on ice with an antibody panel composed of CD64-FITC (22, Beckman Coulter (BC), Miami FL), CD30-PE (BerH83, Beckton-Dickinson (BD), San Jose, CA, RRID:AB_400238), CD5-ECD (BL1a, BC, RRID:AB_3678603), CD40-PE-Cy5.5 (custom conjugate, gift of Jonathan Fromm) or CD40-PerCP-eFluor 710 (1C10, Ebiosciences, San Diego, CA), CD20-PC7 (B9E9, BC), CD15-APC (HI98, BD, RRID:AB_10893192), CD45 APC-H7 (2D1,BD, RRID:AB_1645480) or CD45-Krome Orange (J.33, BC, RRID:AB_2888654), and CD95-Pacific Blue (DX2, Life Technologies, Grand Island, NY), and resuspended in fluorescence-activated cell sorter (FACS) buffer. Cell suspensions were immediately sorted on a special-order FACS Aria research sorter using a 130-micrometer nozzle at 12 psi, separating HRS, B, and T cells from the tumor using 4-way sort settings. Sorted cells were captured in N-2-hydroxyethylpiperazine-N’-2-ethanesulfonic acid buffer solution containing 50% fetal bovine serum. HRS cells and intratumor B cells were used for RNA sequencing.

### Flow cytometry

The proportion of NK cells was examined in 43 CHL tumors and 10 reactive nodes by flow cytometry panel containing antibodies against CD45, CD7, surface CD3 and CD56 (8). NK cells were quantified based on the expression of CD7 and CD56 without surface CD3 among the mononuclear cells. Statistical analysis was performed by a two-tail Student t-test within Excel (Microsoft, Bellevue, WA, RRID:SCR_016137). Expression of CD48 (clone TU145, BD, RRID:AB_396099) on primary HRS cells was examined in 5 cases in the context of a panel containing CD30, CD15, CD20, CD40, CD3, CD95, CD64, and CD45 to identify unrosetted HRS cells as previously described (8).

### RNA Extraction, Library Construction and Next-Generation Sequencing

Cells in catch buffer were carefully pelleted and washed once with phosphate-buffered saline. RNA was extracted using the Arcturus PicoPure RNA Isolation kit (KIT0204, Life Technologies, Carlsbad, CA) with RNAse-Free DNase (79254, Qiagen, Venlo, Netherlands) to isolate high quality RNA from viable flow-sorted cells. RNA was checked for quality and concentration with Bioanalyzer RNA Pico (5067-1513, Agilent, Santa Clara, CA). We applied the SMARTer Ultra Low Input RNA Kit (Clontech, Mountain View, CA) to 1-4 ng of input total mRNA. Illumina-compatible libraries were constructed from cDNA using KAPA LTP Library Preparation Kit (KK8221, Woburn, MA) and sequenced on Illumina HiSeq in paired-end 101-bp mode. Raw data contained 23-71 million read pairs, or a total of 4,600 – 1,400 megabases, per library. The data has been deposited in NCBI GEO, with accession number: GSE301492.

### Differential Gene expression analysis

To determine differential gene expression, raw data were mapped to the human genome GRCh37 using STAR version 2.3.0e. Reads that mapped to any of 52,775 genes annotated in Ensembl(58) were counted using HTSeq-counts v.0.6.1 after which genes without a counts per million reads (CPM) in at least 3 samples were excluded from downstream analysis (59). Count data were normalized using the trimmed mean of M-values (TMM) method and differential gene expression analysis was performed using the limma-voom pipeline (limma version 3.40.6) (60–62). GSEA-4.3.2 was used for Gene set enrichment analysis (GSEA) (63,64). PCA and unsupervised hyerarchichal clustering was performed with samples blinded as to whether they represented HRS of B cells. pheatmap and ggplot2 (version 3.2.1, RRID:SCR_014601) were used to plot the heatmap and cluster-specific trends (65,66). For gene expression comparison among PMBL, ABC-DLBCL and GCB-DLBCL, we used the dataset published by Rosenwald *et al.* (21). The PMBL signature was obtained from Savage *et al*. (20), and gene expression data for PMBL used in this analysis were previously published by Mottok *et al.* (19). The annotated counts and analyses are provided in the Supplementary Datatables and will be submitted to NCBI GEO prior to publication.

### Data Input and Preprocessing

Raw RNA-seq data in FASTQ format was converted into fasta format and then divided into smaller files containing 100,000 reads each, to speed alignment. Each was mapped in parallel on a computational cluster using Basic Local Alignment Search Tool (BLAST, RRID:SCR_004870) (67) to a reference sequence composed of the full set of spliced mRNA transcripts derived from hg19 RefSeq (RRID:SCR_003496). BLAST output was sorted by sequence read id, gene name, and position on the query read, and compressed and formatted on the fly to keep only fields informative for fusions, to reduce the consumption of disk space. Degeneracy on the level of genes was addressed by collapsing hits to multiple different isoforms of the same gene from the same query to one hit per gene per query. Sorted and reduced alignment results were combined back into a single larger file and two mutually exclusive lists were then created: one containing individual reads that mapped exclusively to one gene, and another which contained only reads that mapped uniquely to two or more distinct genes. The former was used to search for split-insert read pairs and the latter for split-reads.

### Virus discovery

RNA-seq files were tested for the presence of viral genomes using the virdetect tool (37). Pre-prepared viral genomes in FASTA format are available for download with the virdetect; these are curated vertebrate viral genomes are obtained from GenBank and then optimized to increase specificity by masking the viral genomes for (a) areas of human homology and (b) areas of low complexity. Following alignment with STAR (with hg38 reference), reads that could not be mapped to the human genome were then mapped to the prepared viral genomes (list of 1893 viruses) by the software.

## ACKNOWLEDGEMENTS

This project was supported by the Department of Pathology and Laboratory Medicine of Weill Cornell Medicine and its Center for Translational Pathology. It was funded in part by NIH grant R01-CA068939 to EC. JR was partially funded by the Tri-I Training Program in Computational Biology and Medicine (5T32GM083937). MR and JR were supported by MSK Cancer Center Support Grant/Core Grant (P30 CA008748).

## AUTHORSHIP CONTRIBUTIONS

EC, MR, and LGR conceived of the experiment, advised on every aspect, conducted validation experiments and wrote the manuscript; MR and JR sorted primary cases, optimized and constructed libraries. JS performed RT-PCR validation. JR, IYK, WD, FW, SZ, BB, and AC analyzed data with LM, OE, and RR. JB, SIP, AEK, MJO, MSL, MJB contributed patient samples. All authors reviewed the manuscript.

