## Supplementary Data for "Transcriptome sequencing of Hodgkin lymphoma Hodgkin and Reed-Sternberg cells reveals escape from NK cell recognition and an unfolded protein response"

**Roshal el al.**

**SUPPLEMENTARY DATA**

**Supplementary Table S1. Patient characteristics**

| Patient # | Age | cHL subtype | Corresponding published case (ref) |
|---|---|---|---|
| 2 | 32 | NS | 2 (1) |
| 3 | 25 | NS | 3 (1) |
| 4 | 58 | CLL+CHL | 4 (1) |
| 5 | 40 | NS | 5 (1) |
| 6 | 51 | NS | 6 (1) |
| 7 | 29 | NS | 7 (1) |
| 8 | 21 | NS | 8 (1) |
| 9 | 76 | MC | 9 (1) |
| 10 | 62 | MC | 10 (1) |
| 11 | 12 | NS | IID_H198432 (2) |
| 12 | 13 | MC | IID_H198438 (2) |
| 13 | 12 | NS | IID_H198431 (2) |
| 14 | 17 | NS | IID_H201345 (2) |
| 15 | 20 | NS | IID_H201358 (2) |
| 16 | 11 | NS | IID_H198451 (2) |
| 17 | N/A | N/A | N/A |
| 18 | 18 | cHL, NOS | N/A |
| 19 | 69 | NS | IID_H198428 (2) |

**Supplementary Table S2. Genes differentially expressed in opposite directions in Roshal et al and Steidl et al.**

| Gene Name | log2FC_Roshal | adj.P.Val_Roshal | log2FC_Steidl | adj.P.Val_Steidl |
|---|---|---|---|---|
| ARHGAP11A | 4.03449127 | 1.11E-07 | -2.722466 | 4.30E-12 |
| CCNA2 | 4.38307052 | 1.36E-08 | -2.8073549 | 5.60E-13 |
| CDKN3 | 4.89290494 | 1.10E-12 | -2.8678965 | 2.00E-13 |
| CENPE | 3.9896341 | 6.23E-09 | -2.3785116 | 9.20E-15 |
| CLU | 6.62875812 | 0.00058648 | -4.7655347 | 8.60E-13 |
| DEPDC1 | 6.41120313 | 1.58E-10 | -2.9448584 | 1.00E-14 |
| E2F8 | 6.93514416 | 4.17E-08 | -2.4059924 | 2.90E-14 |
| FEN1 | 2.93961117 | 9.38E-10 | -2.3504972 | 1.10E-12 |
| HTR3A | -8.0382608 | 2.03E-14 | -3.6667566 | 2.70E-13 |
| ITK | 4.57370943 | 0.000153 | -2.7004397 | 0.00000057 |
| KIF11 | 4.12991817 | 1.22E-08 | -2.4329594 | 3.30E-15 |
| KIF18A | 2.76598644 | 0.00012244 | -2.722466 | 5.50E-12 |
| MAD2L1 | 3.29989895 | 1.74E-11 | -3.169925 | 2.50E-13 |
| MND1 | 3.25072017 | 1.69E-07 | -2.9634741 | 7.10E-14 |
| PANK1 | 4.58122435 | 4.53E-08 | -2.3785116 | 7.50E-10 |
| PHACTR1 | -5.4042134 | 3.87E-09 | 2.48542683 | 0.0000021 |
| PLK4 | 3.62207487 | 5.11E-07 | -2.3504972 | 2.60E-13 |
| SPRED2 | 2.88427631 | 0.00022787 | -3.5484366 | 2.20E-14 |
| TCF4 | -2.6847646 | 1.33E-11 | 3.15380534 | 3.40E-12 |
| TOM1L1 | 5.06489046 | 0.00035108 | -2.4059924 | 4.9E-08 |

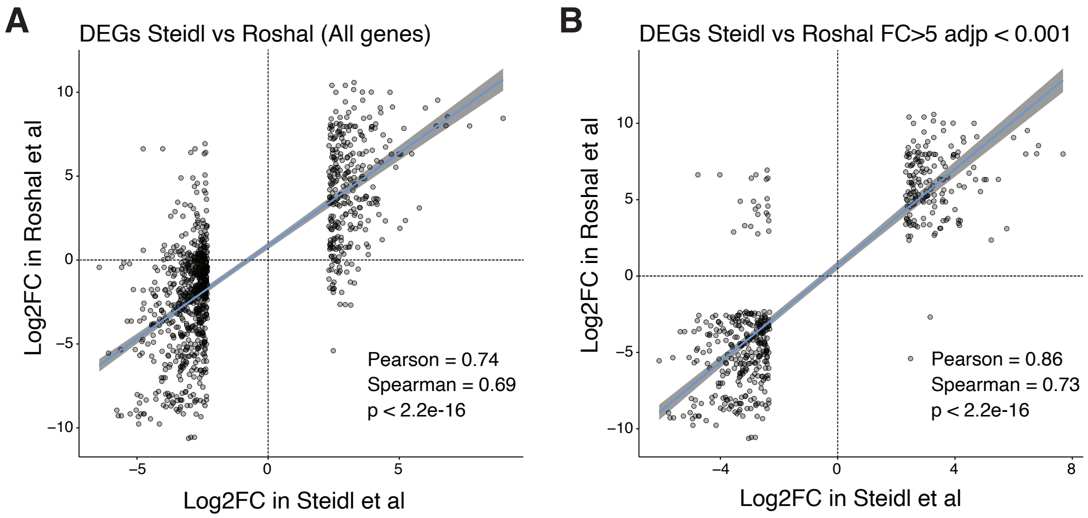


**Supplementary Figure S1. Comparison of gene expression between HRS cells and intra-tumoral B cells using two different approaches.** Gene expression concordance was assessed between RNA sequencing data from flow-cytometry purified HRS cells and previously published data by Steidl et al, which used laser-capture microdissection and Affymetrix GeneChip HG133 V2.0 arrays (3). Fold-change concordance plots are shown for A) all expressed genes, and B) genes meeting the differential expression significance cutoff (FDR<0.001; fold change >5) for HRS cells versus intra-tumoral B cells.

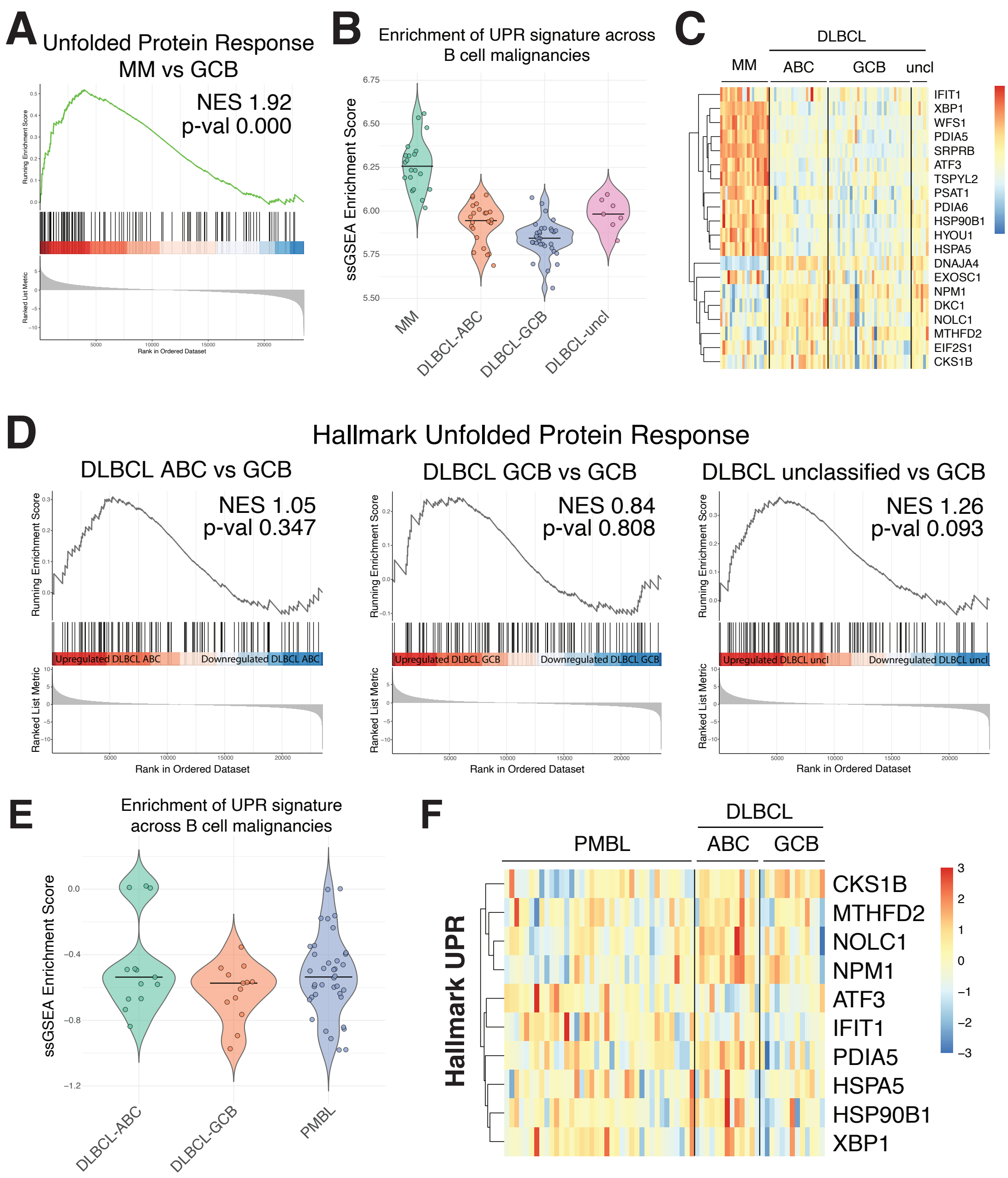


**Supplementary Figure S2. Expression of UPR signature in multiple myeloma and diffuse large B cell lymphoma.** (**A**) Gene set enrichment analysis (GSEA) for the Hallmark of Unfolded Protein Responses (UPR) was performed on multiple myeloma (MM) vs germinal centre B cells (GCB). (**B**) Enrichment of UPR signatures across MM and different sub-classifications of diffuse large B cell lymphoma (DLBCL), using expression data from Agirre et al. (4). (**C**) Heatmaps of genes associated with Hallmark of unfolded protein response (UPR) pathway as shown in Figure 2F in MM and different sub-classifications of diffuse large B cell lymphoma (DLBCL) in panel B. (**D**)GSEA for the Hallmark of Unfolded Protein Responses (UPR) analysis for DLBCL samples separated to performed on multiple myeloma (MM) vs germinal centre B cells (GCB). (**E**) Enrichment of UPR signatures in PMBL samples as published by Rosenwald et al. (**F**) Heatmaps of genes associated with Hallmark of unfolded protein response (UPR) pathway as shown in Figure 2F in PMBL samples, as published by Rosenwald et al. (5) ABC-DLBCL=Activated B-cell-like diffuse large B cell lymphoma; GCB-DLBCL=germinal center B-cell-like diffuse large B cell lymphoma; uncl-DLCBL=unclassified-DLBCL. Colors in B and D represents relative expression: red indicates higher and blue indicates lower expression relative to the gene's mean.

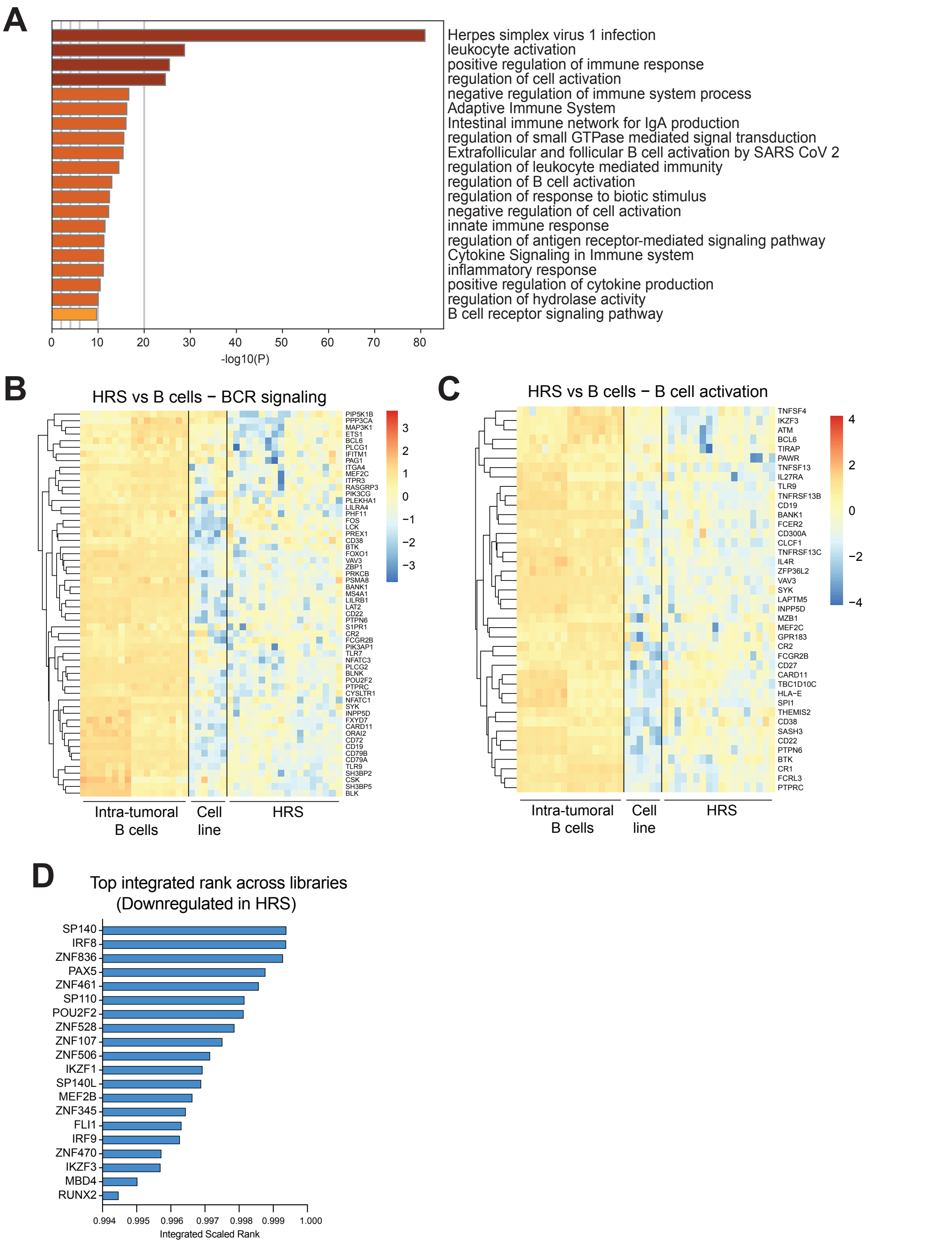


**Supplementary Figure S3. Downregulation of B cell signalling and activation pathways in HRS cells. (A)** Gene ontology analysis of genes downregulated in HRS compared to intra-tumoral B cells (adjp≤0.05; logFC≥ 1 or logFC ≤ -1). Heatmaps for (**B**) B cell receptor (BCR) signalling and (**C**) B cell activation genes differentially regulated between HRS and intra-tumoral B cells (adjp≤0.05; logFC≥ 1 or logFC ≤ -1). (**D**) Transcription factor target over-representation analysis was conducted using ChIP-X Enrichment Analysis version 3 (ChEA3) for genes downregulated in HRS relative to intra-tumoral B cells (adj-p≤0.001, logFC≤-2.32). Colors in B and C represents relative expression: red indicates higher and blue indicates lower expression relative to the gene's mean.

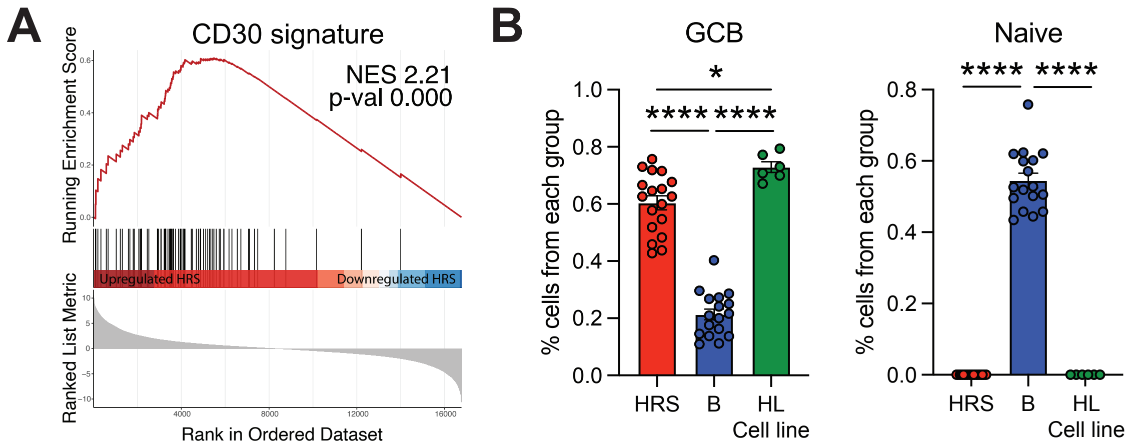


**Supplementary Figure S4. Comparison of HRS cell signature with other lymphoma subsets.** (**A**) Gene set enrichment analysis (GSEA) for genes upregulated in CD30+ cells (Weniger et al) comparing HRS to intra-tumoral B cells. (**B**) Estimated proportion of germinal centre B cells (GCBs) and naïve B cells in HRS, intra-tumoral B cells and cell line samples.

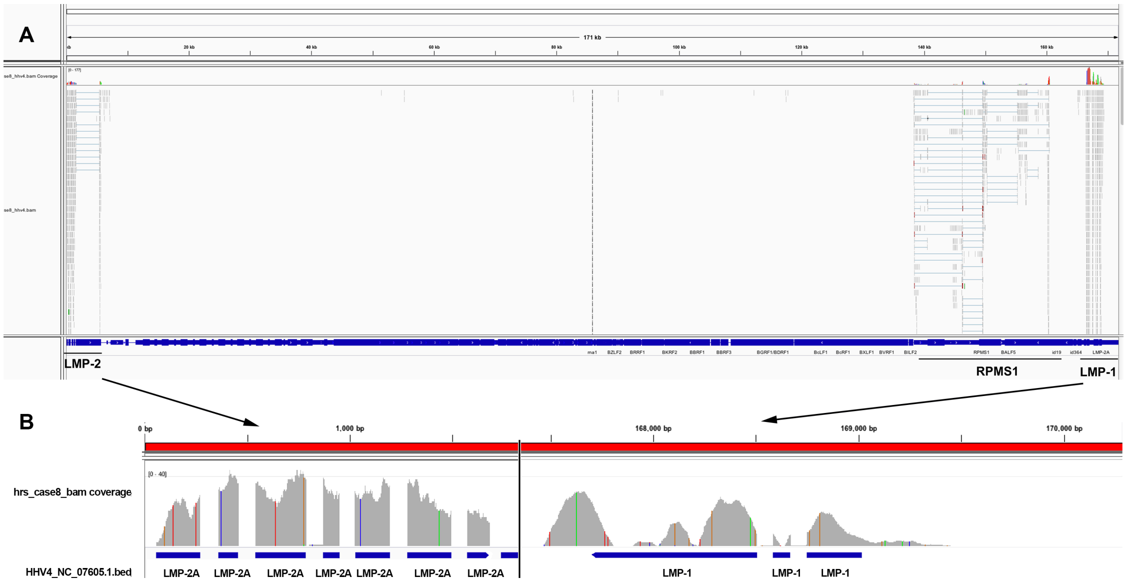


**Supplementary Figure S5. Detection of EBV in cHL.** Alignment of transcripts from Case 8 with a reference EBV genome using Integrative Genomics Viewer (IGV) is shown. (A) The entire genome shows most reads aligning to the regions surrounding the terminal repeats, which are at the beginning and end of the linear genome, where LMP-1 and LMP1 are located, although some reads were found in the RPMS1 region as well. (B) A higher scale view of the coverage in the LMP-2A and LMP1 genes shows expression of these genes is provided.
